# SARS-CoV-2 spike glycoprotein-reactive T cells can be readily expanded from COVID-19 vaccinated donors

**DOI:** 10.1101/2021.05.27.446089

**Authors:** Pavla Taborska, Jan Lastovicka, Dmitry Stakheev, Zuzana Strizova, Jirina Bartunkova, Daniel Smrz

**Author notes:** Address correspondence to Dr. Daniel Smrz: Department of Immunology, Second Faculty of Medicine, Charles University and University Hospital Motol, V Uvalu 84, 150 06 Praha 5, Czech Republic.

## Abstract

**Introduction:** The COVID-19 vaccine was designed to provide protection against infection by the severe respiratory coronavirus 2 (SARS-CoV-2) and coronavirus disease 2019 (COVID-19). However, the vaccine’s efficacy can be compromised in patients with immunodeficiencies or the vaccine-induced immunoprotection suppressed by other comorbidity treatments, such as chemotherapy or immunotherapy. To enhance the protective role of the COVID-19 vaccine, we have investigated a combination of the COVID-19 vaccination with *ex vivo* enrichment and large-scale expansion of SARS-CoV-2 spike glycoprotein-reactive CD4^+^ and CD8^+^ T cells.

**Methods:** SARS-CoV-2-unexposed donors were vaccinated with two doses of the BNT162b2 SARS-CoV-2 vaccine. The peripheral blood mononuclear cells of the vaccinated donors were cell culture-enriched with T cells reactive to peptides derived from SARS-CoV-2 spike glycoprotein. The enriched cell cultures were large-scale expanded using the rapid expansion protocol (REP) and the peptide-reactive T cells evaluated.

**Results:** We show that vaccination with the SARS-CoV-2 spike glycoprotein-based mRNA COVID-19 vaccine induced humoral response against SARS-CoV-2 spike glycoprotein in all tested healthy SARS-CoV-2-unexposed donors. This humoral response was found to correlate with the ability of the donors’ PBMCs to become enriched with SARS-CoV-2 spike glycoprotein-reactive CD4^+^ and CD8^+^ T cells. Using an 11-day rapid expansion protocol, the enriched cell cultures were expanded nearly a thousand fold, and the proportions of the SARS-CoV-2 spike glycoprotein-reactive T cells increased.

**Conclusions:** These findings show for the first time that the combination of the COVID-19 vaccination and *ex vivo* T cell large-scale expansion of SARS-CoV-2-reactive T cells could be a powerful tool for developing T cell-based adoptive cellular immunotherapy of COVID-19.

## Introduction

COVID-19 is transforming into more severe and contagious forms as the severe respiratory coronavirus 2 (SARS-CoV-2) mutates during the pandemic [1]. The recently appeared new mutations of the virus seem to evade the immune system more efficiently, including evasion of the coronavirus disease 2019 (COVID-19) vaccine-induced immunity [2]. This evasion lies in the decreased capability of SARS-CoV-2-specific antibodies to neutralize the virus efficiently [3]. Since the antibody-mediated protection depends on the structure of the target antigen, any mutation causing a productive conformational change of the target antigen can decrease the antibody binding and erode its protective role [4, 5]. The current COVID-19 vaccines are nearly exclusively targeting a single protein of the virus, the spike glycoprotein, so chances of evasion could be high.

The antiviral immune response also relies on adaptive cellular immunity where the antiviral effectors are, instead of antibodies, the cytotoxic CD8^+^ T cells which recognize infected cells expressing viral proteins [6]. Unlike antibodies, the viral proteins are recognized in the form of protein fragments (peptides) presented in the context of the major-histocompatibility complexes and T cell receptors (TCR) [6]. Recent studies show that SARS-CoV-2 T cell-based immunity is negligibly impacted by the current mutated variants of SARS-CoV-2 [7] and, therefore, could counteract the debilitating impact these mutations might have on the parallel humoral immunity [8]. However, many immunocompromised patients, patients with immunodeficiencies, or patients with a comorbidity treatment-suppressed immunity, such as patients undergoing chemotherapy or immunotherapy, may not sufficiently mobilize the cellular immunity against SARS-CoV-2 after vaccination. It is, therefore, necessary to find new ways to enhance their cellular immunity against the virus.

This study examined the impact of the COVID-19 vaccine on the induction of the humoral and cellular responses in 8 healthy SARS-CoV-2-unexposed donors. We investigated whether the cellular response to the vaccine could be enhanced by the *ex vivo* enrichment and large-scale expansion and hence represent an avenue for promoting the SARS-CoV-2-specific cellular immunity in patients who could not fully benefit from the COVID-19 vaccines.

## Results

### COVID-19 vaccination induces SARS-CoV-2 spike glycoprotein-specific antibodies

We first investigated the humoral response of the mRNA SARS-CoV-2 spike glycoprotein-based COVID-19 vaccine, BNT162b2, in 8 healthy donors who tested negative for the presence of antibodies specific to SARS-CoV-2 spike glycoprotein and who reported no previous history of COVID-19 and/or positive tests for SARS-CoV-2. The donors were vaccinated with two doses of the vaccine within a (3–4)-week interval. Samples, the serum and unclotted blood, were collected during the 2 days before each vaccination and 3–4 weeks after the second dose of the vaccine. Using a microblot system, the sera were analyzed for the presence of SARS-CoV-2-specific IgA, IgG, or IgM antibodies against the virus proteins: receptor-binding domain (RBD) and S2 subunits of the spike glycoprotein (S2), nucleocapsid protein (NCP), envelope protein (EP), and papain-like protease (PLP) [9]. As negative controls, the sera were analyzed for the presence of antibodies specific to human ACE-2 protein [10] or proteins from other coronaviruses: S1 subunit of the Middle East respiratory syndrome-related coronavirus spike glycoprotein (MERS-CoV) [11], nucleocapsid protein of SARS-CoV [12], human coronavirus 229E [13] and NL63 [14]. As shown in Fig. 1A, the COVID-19 vaccination induced no production of SARS-CoV-2 unrelated antibodies. The COVID-19 vaccination also did not induce production of antibodies specific to the SARS-CoV-2 nucleocapsid protein, envelope protein, or papain-like protease (Fig. 1B), which indicated no previous SARS-CoV-2 infection. On the other hand, the vaccine induced the production of antibodies specific to SARS-CoV-2 glycoprotein. As shown in Fig. 1B (middle panel), the RBD-specific IgG antibodies were already induced in all the tested donors after the first dose of the vaccine and their levels were further enhanced after the second dose of the vaccine. Only one donor after the first dose and two donors after the second dose of the vaccine produced IgG antibodies against the S2 subunit of the virus spike glycoprotein (Fig. 1B, middle panels), indicating stronger immunogenicity of its RBD domain. This stronger immunogenicity was even more pronounced for the IgA-specific antibodies where the vaccine induced the production of only the RBD-specific IgA antibodies (Fig. 1B, left panels). The production of these IgA antibodies was determined only in 7 donors because one donor in the cohort was diagnosed IgA deficient. The production of IgM antibodies was detected only in 1 donor and only after the first dose of the vaccine. The microblot results showed that the COVID-19 vaccination predominantly induced a specific IgA or IgG antibody response against the RBD of the SARS-CoV-2 spike glycoprotein and less frequently a specific IgG antibody response against the S2 subunit of the SARS-CoV-2 spike glycoprotein.

**Figure 1.**
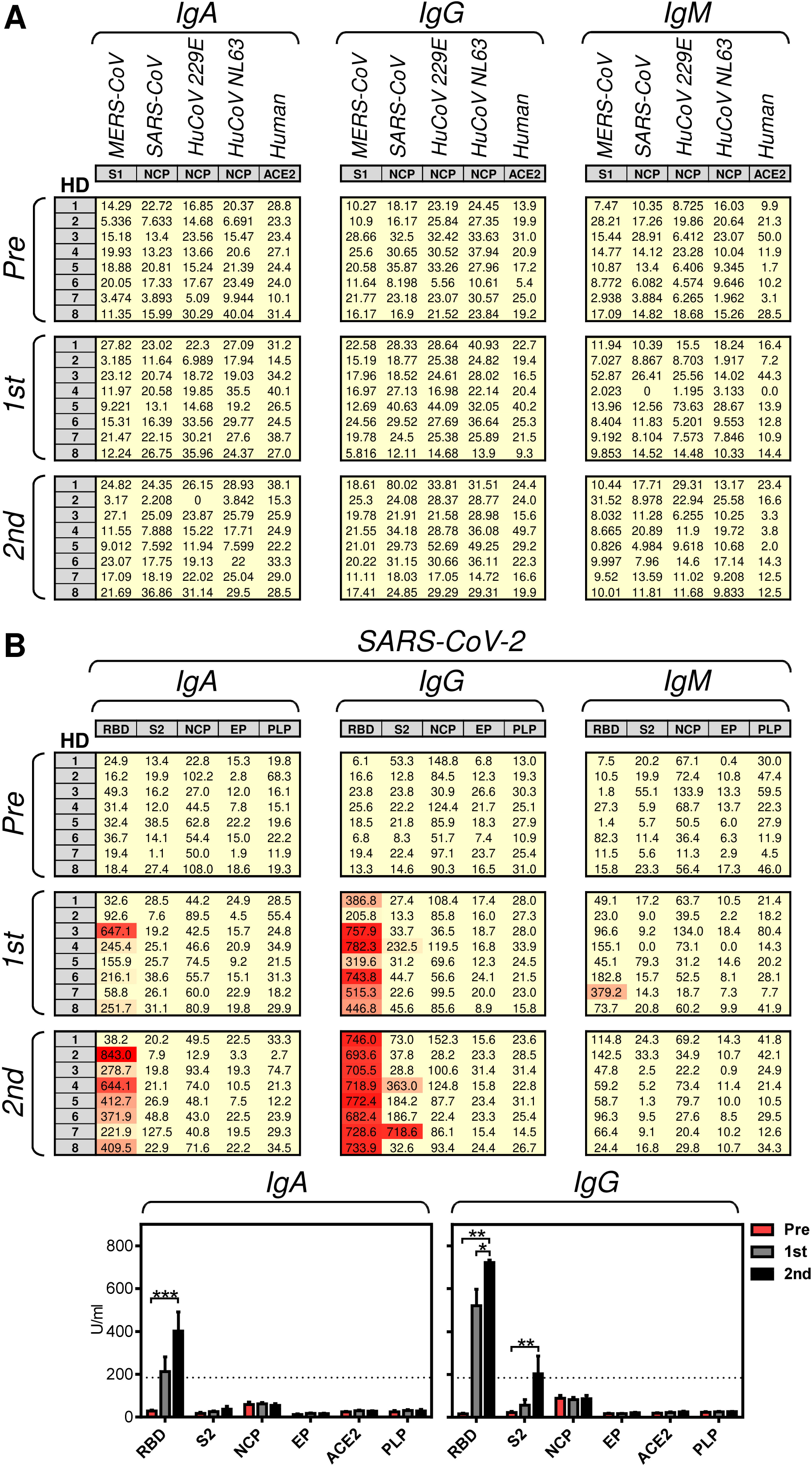
The serum levels of the antigen-specific IgG, IgA, and IgM antibodies determined by the Microblot-Array COVID-19. (A–B) The 2 vaccine doses were administered to 8 healthy donors (HD) with 3 to 4 weeks between the doses. The serum samples were collected 0–2 days before each vaccine dose (Pre, 1st), and 3–4 weeks after the second vaccine dose (2nd). In A, the serum levels (U/ml) of IgA, IgG, and IgM antibodies specific to non-SARS-CoV-2 proteins; Middle East respiratory syndrome-related coronavirus spike glycoprotein S1 subunit (MERS-CoV, S1), SARS-CoV nucleocapsid protein (SARS-CoV, NCP), human coronavirus 229E NCP (HuCoV 229E, NCP), human angiotensin-converting enzyme (Human, ACE-2). In B, the serum levels (U/ml) of IgA, IgG, and IgM antibodies specific to SARS-CoV-2 proteins; spike glycoprotein receptor-binding domain (RBD) and S2 domain (S2), nucleocapsid protein (NCP), E protein (EP), and papain-like protease (PLP). (C) The data in A and B were evaluated. The bars represent mean of values±SEM and significances of differences among the groups (Pre, 1st, 2nd) for individual proteins are indicated (\**P*<0.05, \*\**P*<0.01, \*\*\**P*<0.001, IgA: *n* = 7 (HD1 was excluded because diagnosed as IgA deficient) and IgG: *n* = 8 healthy donors (HD), 1-way ANOVA with the Dunn’s posttest).

### COVID-19 vaccination induces SARS-CoV-2 spike glycoproteinspecific cellular immunity

The cellular immunity against SARS-CoV-2 is increasingly considered to be as important for the effective protection against the virus as the humoral immunity [15]. Since our data showed that the COVID-19 vaccine specifically induced humoral response against SARS-CoV-2 spike glycoprotein, we next investigated whether the COVID-19 vaccination impacted the reactivity of the donors’ CD4^+^ and CD8^+^ T cells to peptides derived from SARS-CoV-2 spike glycoprotein. We first found that the COVID-19 vaccination did not affect the viability of the isolated donors’ peripheral blood mononuclear cells (PBMCs) (Fig. 2A and 2B). The vaccination also had a minimal effect on the proportions of T cells and their CD4^+^ and CD8^+^ subpopulations (Fig. 2A and 2B). To determine the reactivity of the donors’ CD4^+^ and CD8^+^ T cells to peptides derived from the SARS-CoV-2 spike glycoprotein, the donors’ PBMCs were stimulated with a pool of peptides derived from the glycoprotein (Fig. 2C). The peptide pool-stimulated cells were either analyzed by intracellular cytokine staining (ICS) after a 5 h stimulation or cultured for 12 days in the presence of IL-2 to enrich the cell cultures for the peptide-specific T cells [16]. Following the stimulation of the 12-day-enriched cell cultures with the peptide pool for 5 h, the presence of the peptide-specific T cells was determined by ICS (Fig. 2C). As shown, the 12-day cell culture enrichment decreased the viability of the cultured cells (Fig. 2D, left panels) but increased the content of T (CD3^+^) cells in the samples obtained after the COVID-19 vaccinations (Fig. 2D, middle panels). The cell culture enrichment also altered the proportions of CD4^+^ and CD8^+^ populations of T cells in samples obtained after the first dose of the COVID-19 vaccine (Fig. 2D, two right hand panels in the second row).

**Figure 2.**
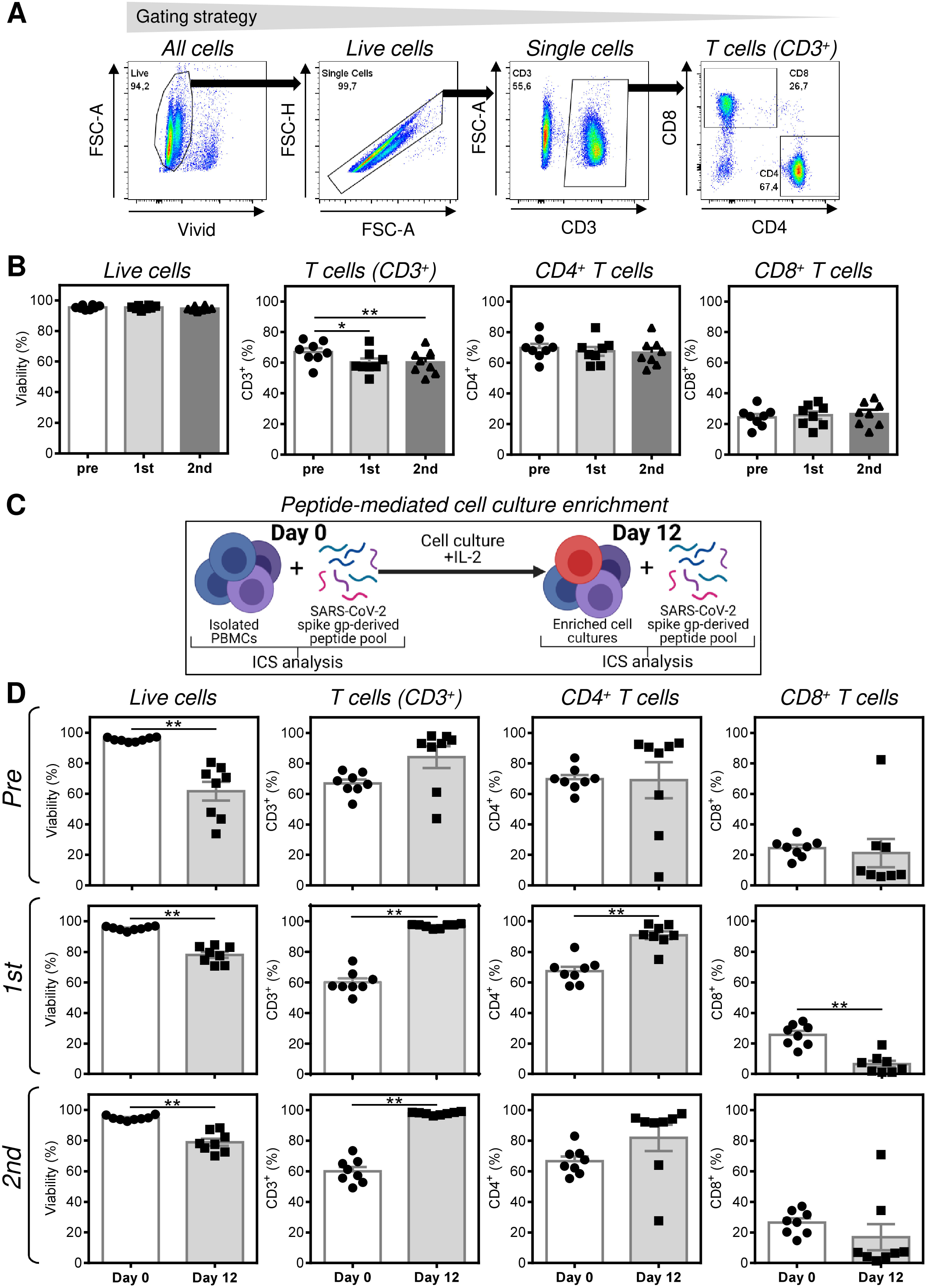
Characterization of the isolated PBMCs and peptide-enriched cell cultures. (A–B) Isolated PBMCs from samples obtained as in Fig. 1 before each vaccine dose (Pre, 1st), and 3–4 weeks after the second vaccine dose (2nd) were characterized by flow cytometry. In A, the gating strategy of flow cytometry data. In B, the cells were analyzed for their viability, the proportion of T cells (CD3^+^), and proportions of CD4^+^ and CD8^+^ populations of T cells. (C) Schematic presentation of the peptide-mediated enrichment and intracellular cytokine staining (ICS) analyses. (D) PBMCs in A–B were peptide-enriched for 12 day and analyzed as in A–B. Their viability (Vivid^-^), the proportion of T cells (CD3^+^), and proportions of CD4^+^ and CD8^+^ populations of T cells of the 12-day peptide-enriched cultures (Day 12) were determined, and the data compared with PBMCs before the enrichment (Day 0). In B and D, the bars represent mean of values±SEM and significances of differences between PBMCs (Day 0) and the peptide-enriched cell cultures (Day 12) were determined for individual groups of the collected samples (Pre, 1st, 2nd; \*\**P*<0.01, *n* = 8 healthy donors, Wilcoxon matched-pairs signed-ranks test).

The presence of peptide-specific T cell populations was determined by ICS of TNFα- and IFNγ-producing T cells (Fig. 3A). As shown in Fig. 3B, the isolated PBMCs from all donors and regardless of the COVID-19 vaccination contained no detectable TNFα-, IFNγ- or TNFα/IFNγ-producing CD4^+^ or CD8^+^ T cell populations reactive to the SARS-CoV-2 spike glycoprotein-derived peptides. However, the 12-day peptide-mediated enrichment significantly enriched cell cultures with the peptide-reactive T cell populations (Fig. 3C). As shown, the peptide-enriched cell cultures already contained TNFα-producing CD4^+^ T cell population reactive to the peptides, and this population was significantly higher in cell samples enriched after the second dose of the vaccine than in cell samples enriched before the vaccination (Fig. 3C, top left panel). No effect of the vaccination on the enrichment with the peptide-reactive IFNγ- or TNFα/IFNγ-producing CD4^+^ T cells was observed because no such populations were detected in the 12-day-enriched cell cultures (Fig. 3C, middle and right top panels). However, the vaccination had a robust impact on the enrichment of cell cultures with the peptide-reactive CD8^+^ T cells. As shown in Fig. 3C (bottom panels), the pre-vaccination samples were not enriched with the peptide-reactive CD8^+^ T cells, showing that the tested SARS-CoV-2-unexposed donors failed to attain a peptide-mediated *ex vivo* enrichment with the SARS-CoV-2 spike glycoprotein-reactive CD8^+^ T cells. This failure was overcome by the COVID-19 vaccination because after the second dose of the vaccine, the cell cultures become significantly enriched with the peptide-reactive CD8^+^ T cells (Fig. 3C, bottom panels). Moreover, this reactivity was shown not only for the TNFα-producing CD8^+^ T cells (Fig. 3C, bottom left panel) but also for the IFNγ- or TNFα/IFNγ-producing CD8^+^ T cells (Fig. 3C, middle and left bottom panels). These data showed that the COVID-19 vaccination significantly promoted the ability of the donors’ PBMCs to become *ex vivo* enriched with the SARS-CoV-2 spike glycoprotein-reactive CD4^+^ and CD8^+^ T cells.

**Figure 3.**
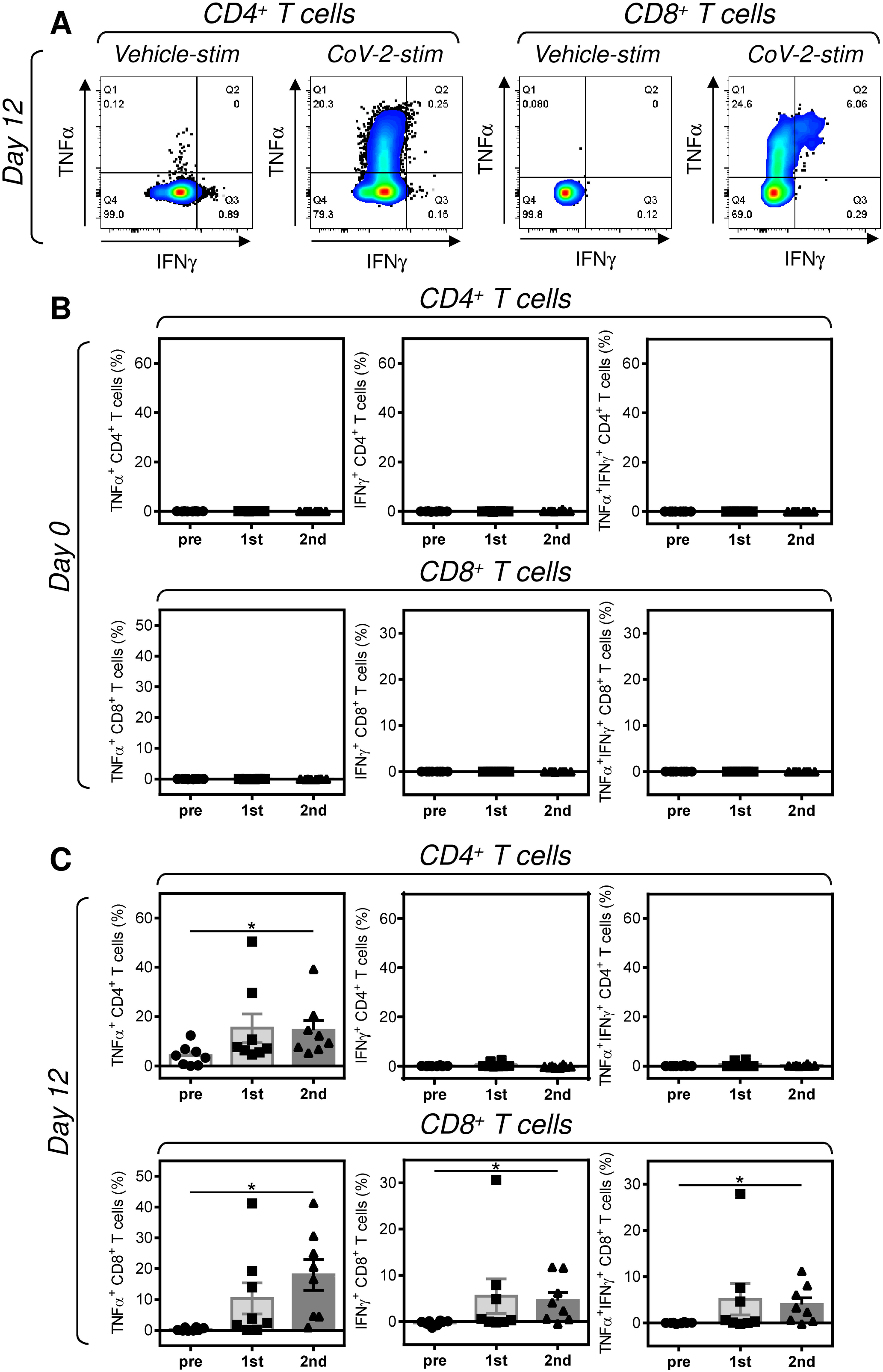
Reactivity of the isolated PBMCs and peptide-enriched cell cultures to SARS-CoV-2 spike glycoprotein-derived peptides. (A) The isolated PBMCs (Day 0) and peptide-enriched cell cultures (Day 12) were (CoV-2-stim) or were not (Vehicle-stim) stimulated with SARS-CoV-2 spike glycoprotein-derived peptides and the proportions of TNFα-, IFNγ-, or TNFα/IFNγ-producing CD4^+^ and CD8^+^ T cells determined by intracellular cytokine staining (ICS). The gating strategy of the flow cytometry data. (B–C) The proportions of reactive T cells in PBMCs (Day 0) (B) and the 12-day peptide-enriched cell cultures (Day 12) (C) were calculated as the difference between the proportions of the cytokine-producing T cells of the vehicle-stimulated sample and the peptide-stimulated sample of the same donor. In B and C, the bars represent mean of values±SEM and significances of differences among the groups (Pre, 1st, 2nd) for TNFα-, IFNγ-, or TNFα/IFNγ-producing CD4^+^ and CD8^+^ T cells are indicated (\**P*<0.05, *n* = 8 healthy donors, 1-way ANOVA with the Dunn’s posttest).

### COVID-19 vaccination-induced humoral response correlates with the SARS-CoV-2 spike glycoprotein peptide reactivity of the peptide-enriched PBMCs

Our data showed that the COVID-19 vaccine promoted both humoral and cellular immunity against SARS-CoV-2 spike glycoprotein. We next analyzed whether the extent of the humoral response correlated with the ability of PBMCs to become enriched with SARS-CoV-2 spike glycoprotein-reactive CD4^+^ and CD8^+^ T cells. As shown in Fig. 4, the extent of the humoral response correlated with the ability of PBMCs to become enriched with SARS-CoV-2 spike glycoprotein-reactive CD4^+^ and CD8^+^ T cells. The levels of the RBD-specific IgG antibodies were found to correlate with the extent of the PBMCs’ *ex vivo* enrichment with the SARS-CoV-2 spike glycoprotein-reactive TNFα-producing CD4^+^ and TNFα-, IFNγ- or TNFα/IFNγ-producing CD8^+^ T cells (Fig. 4A). Comparable data were obtained upon the correlations with the levels of RBD-specific IgA antibodies (Fig. 4A). The only exception was with no correlation found for the TNFα-producing CD4^+^ T cells (Fig. 4B, left panel). Overall, the data showed a close association between the COVID-19 vaccination-induced humoral and cellular responses against the SARS-CoV-2 spike glycoprotein.

**Figure 4.**
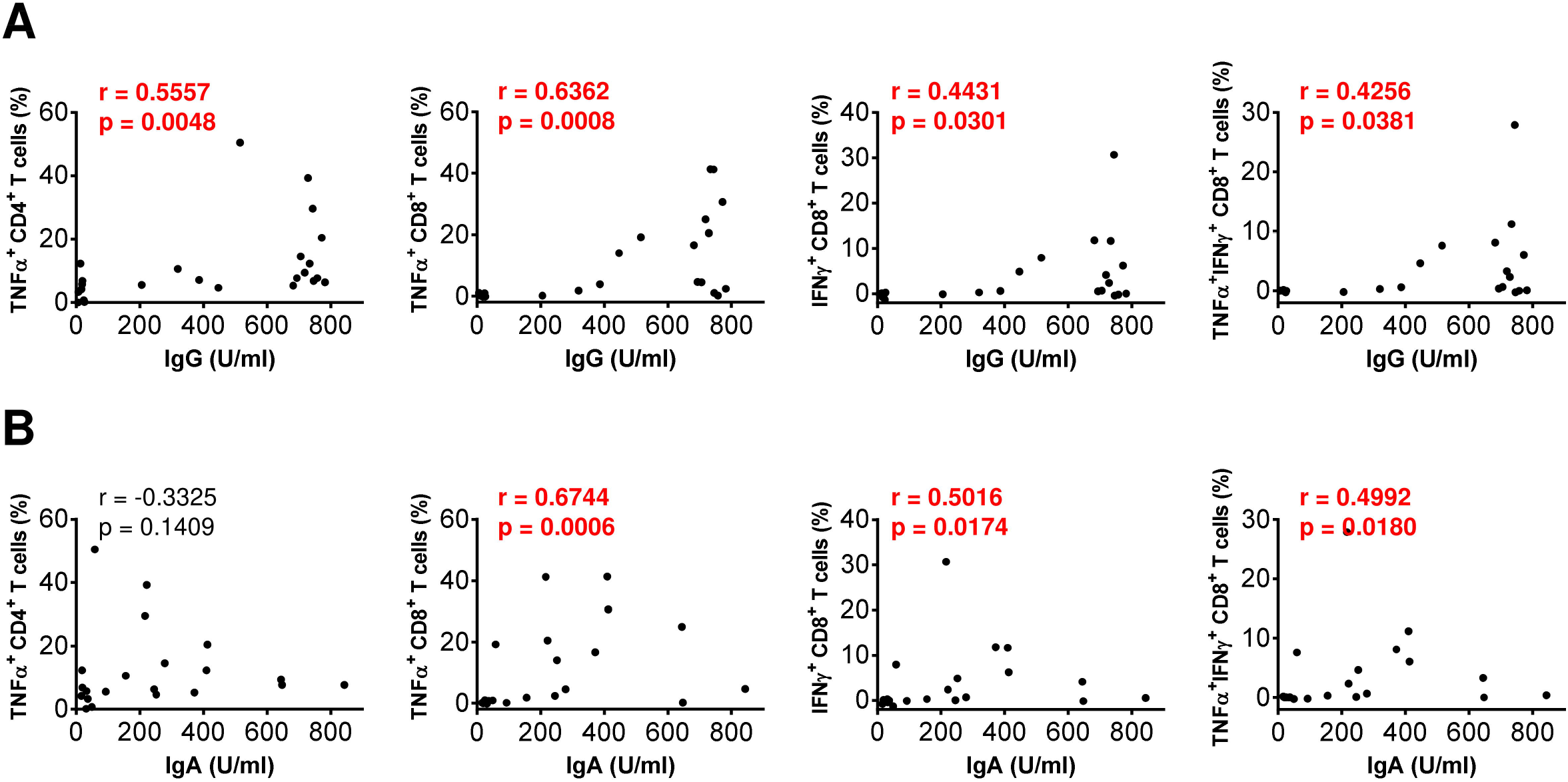
The association between the humoral response and T cell reactivity during the vaccination. (A–B) The correlations between the levels (U/ml) of SARS-CoV-2 spike glycoprotein receptor-binding domain (RBD)-specific antibodies and the proportions of SARS-CoV-2 spike glycoprotein receptor-reactive TNFα-producing CD4^+^ and TNFα-, IFNγ-, or TNFα/IFNγ-producing CD8^+^ T cells were evaluated by Spearman’s rank-order correlation coefficient (r) and the significance (*P* value) determined (IgA: *n* = 7 (HD1 was excluded because diagnosed as IgA deficient) and IgG: *n* = 8 healthy donors).

### The number *of ex vivo*-enriched SARS-CoV-2 spike glycoprotein-reactive CD4^+^ and CD8^+^ T cells can be large-scale expanded in the cell culture

The peptide-enrichment experiments showed that COVID-19 vaccination could significantly enhance or even induce the PBMC’s ability to become enriched with SARS-CoV-2 spike glycoprotein-reactive CD4^+^ and CD8^+^ T cells. We further investigated whether this enrichment could also have the potential to become an avenue for a T cell-based immunotherapy of COVID-19. We used the peptide-enriched cell cultures from the donors’ PBMCs after the second dose of the vaccine and expanded the number of cells using the rapid expansion protocol (REP) [17] (Fig. 5A). As shown in Fig. 5B, the 11-day REP led to a 743.6 [range from 566.3 to 912.0, n = 8, 95% CI 632.0–855.2] cell number fold increase. The expanded cells were of higher viability, with increased proportions of T cells and similar proportions of CD4^+^ and CD8^+^ T cell populations (Fig. 5C–5F). Importantly, the expanded cell cultures became further enriched with the peptide-specific CD4^+^ and CD8^+^ T cells (Fig. 5G). As shown in Fig. 5G, the enrichment was significant for both the peptide-reactive CD4^+^ T cells producing TNFα-, IFNγ- or TNFα/IFNγ and CD8^+^ T cells producing TNFα. The findings showed that the combination of COVID-19 vaccination, peptide-mediated enrichment, and REP could lead to the production of therapeutically relevant numbers of SARS-CoV-2 spike glycoprotein-reactive CD4^+^ and CD8^+^ T cells.

**Figure 5.**
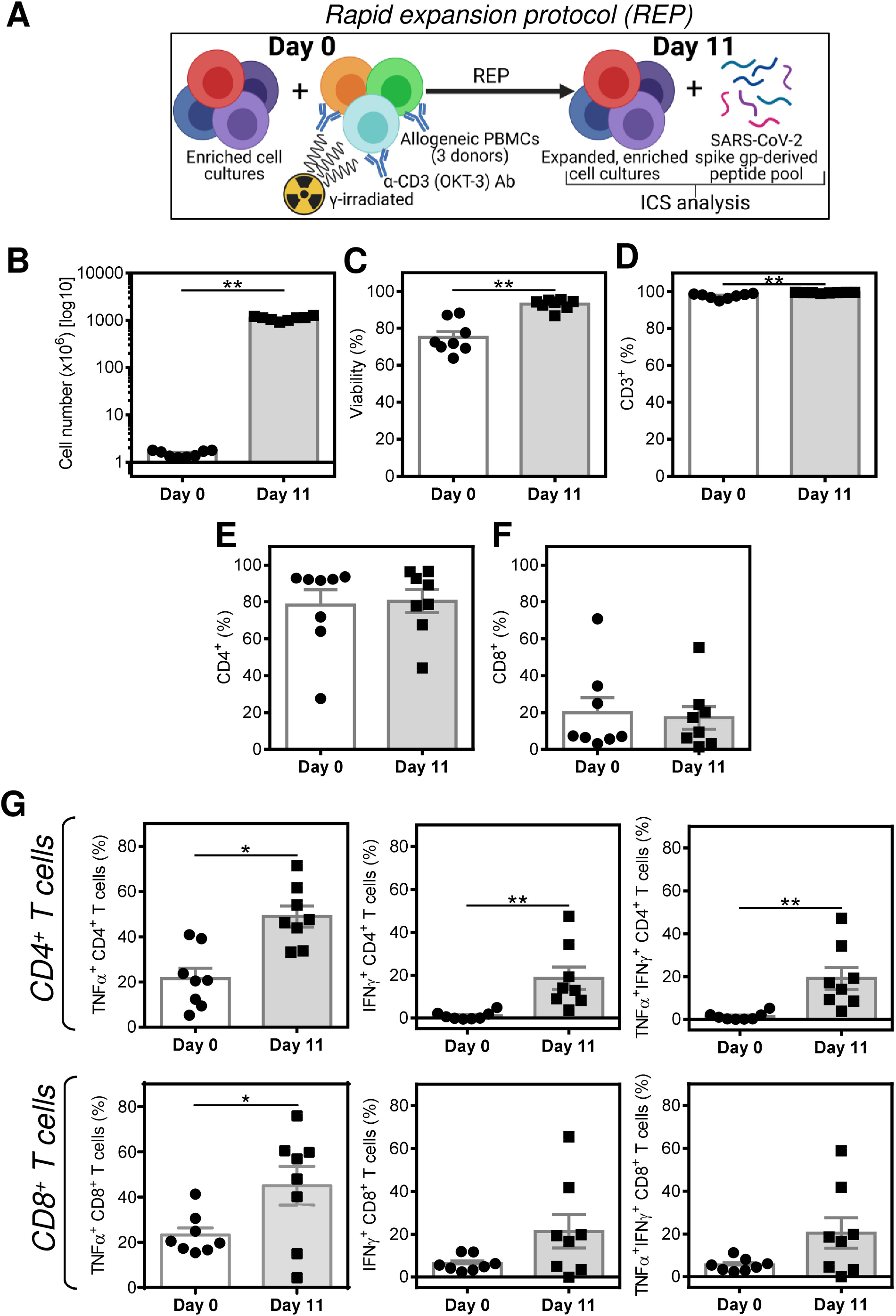
Large-scale expansion of the peptide-enriched cell cultures from the 2nd dose-vaccinated donors using rapid expansion protocol (REP). (A) Schematic presentation of REP and the intracellular cytokine staining (ICS) analysis. (B–F) Cell numbers before (Day 0) and after (Day 11) the REP (B), their viability (C), the proportion of T cells (CD3^+^) (D), and proportions of CD4^+^ (E) and CD8^+^ (F) T cell populations. (G) The peptide-enriched (Day 0) and their REP-expanded counterpart (Day 11) were, or not, stimulated with SARS-CoV-2 spike glycoprotein-derived peptides and the proportions of TNFα-, IFNγ-, or TNFα/IFNγ-producing CD4^+^ and CD8^+^ T cells determined by intracellular cytokine staining (ICS). The proportions of reactive T cells were calculated as the difference between the proportions of the cytokine-producing T cells of the vehicle-stimulated sample and the peptide-stimulated sample of the same donor. In B–G, the bars represent mean of values±SEM and significances of differences among the groups (Day 0, Day 11) for TNFα-, IFNγ-, or TNFα/IFNγ-producing CD4^+^ and CD8^+^ T cells are indicated (\**P*<0.05, \*\**P*<0.01, *n* = 8 2nd dose-vaccinated healthy donors, Wilcoxon matched-pairs signed-ranks test).

## Discussion

This study showed that COVID-19 vaccines could elicit both a humoral and cellular response against the virus. Using the cell culture techniques and peptides derived from the virus antigen, the vaccine-induced antigen-reactive T cells can be *ex vivo*-enriched and large-scale-expanded and as such represent a potential therapeutic tool for the enhancement of cellular immunity after COVID-19 vaccination.

The previous reports have shown that the BNT162b2 vaccine potentiated both the humoral and cellular responses [18]. Our data confirmed that vaccination of healthy donors with this vaccine indeed induced a humoral immune response that led to the production of SARS-CoV-2 spike glycoprotein-specific IgG and IgA antibodies. This response was highly specific as the vaccination induced no detectable antibodies specific to other SARS-CoV-2 proteins or proteins from other coronaviruses. These data, therefore, confirmed the precision of the vaccine-based prophylactic immunotherapy.

The cellular immunity is the important layer of the immune protection against viruses as it prevents the virus amplification after infection [19, 20]. The effector cells of this arm of immunity are primarily the cytotoxic CD8^+^ T cells [6]. This study showed that no such SARS-CoV-2 spike glycoprotein-specific CD8^+^ T cells were detected in the peripheral blood of either non-vaccinated or 2-dose-vaccinated donors. These cells were also not detectable in the non-vaccinated donors even after the peptide-mediated enrichment. But, once the donors obtained 2 doses of the COVID-19 vaccine, the peptide-mediated enrichment already produced cell cultures containing the SARS-CoV-2 spike glycoprotein-specific CD8^+^ T cells. These data showed that the vaccination was important for increasing the frequency of the SARS-CoV-2 spike glycoprotein-specific CD8^+^ T cells to the levels that allow their peptide-mediated enrichment in the cell culture. These findings corroborate previous reports showing increased frequencies of T cells reactive to peptides derived from the tumor-associated antigens in the peptide-enriched cell cultures after the patients’ vaccination with *ex vivo*-produced dendritic cells loaded with whole inactivated tumor cells [21, 22].

Our results showed that humoral and T cell-based immune responses went hand in hand in the tested healthy donors. However, patients with compromised immunity or undergoing therapies that compromise their immunity may not respond well with both arms of the adaptive immunity, and the protective potential of the COVID-19 vaccines can then be undermined in these patients [23]. The large-scale expanded antigenspecific T cells have been utilized for adoptive cellular immunotherapy (ACI) of cancer [24]. Both prophylactic and therapeutic anti-viral ACI has also been utilized after the hematopoietic stem cell (HSC) transplantations, where viral infections are an important cause of morbidity and mortality [25, 26]. The restoration of the viral immunity is often successfully attained by adoptive transfer of the HSC donor’s *ex vivo* expanded virus-specific CD8^+^ T cells [25, 26]. The expanded SARS-CoV-2-reactive T cells could, therefore, be also implemented in these therapeutic strategies to compensate for insufficiencies of the SARS-CoV-2-specific cellular immunity. The findings of the study show that the combination of the COVID-19 vaccines with the *ex vivo* peptide-mediated enrichment and large-scale expansion could represent a viable approach for the production of T cells for cellular therapy of COVID-19.

## Materials and methods

### Donors and COVID-19 vaccination

The study involved 8 healthy donors who were negative for SARS-CoV-2 spike glycoprotein-specific antibodies and who reported no previous history of COVID-19 or positivity for SARS-CoV-2 infection. The median age of the donors was 46.0 years (range 32–72 years) and the samples were obtained between October 2020 and February 2021. The donors were vaccinated with the BNT162b2 SARS-CoV-2 vaccine (Pfizer-BioNTech) with two doses with a 3 to 4 week interval between each dose. The donors’ samples, the peripheral blood serum and unclotted peripheral blood, were collected up to 2 days before the first and second dose of the vaccine and 3–4 weeks after the second dose of the vaccine. The serum was separated by centrifugation at 3000 rpm for 5 min at room temperature and cryopreserved. PBMCs from the unclotted peripheral blood were isolated as previously described [27] and cryopreserved [RPMI 1640 medium (Thermo Scientific, Waltham, MA), 10% human plasma serum (One Lambda, Canoga Park, CA), 10% DMSO (Sigma-Aldrich, St. Louis, MO, USA), 100 U/ml penicillin-streptomycin, and 2 mM GlutaMax (Thermo Scientific)]. Each donor provided signed written informed consent for the use of their blood-derived products for future research and all experimental protocols were approved by the ethical standards of the institutional research committee – the Ethics Committee of the University Hospital Motol in Prague, and performed in accordance with the 1964 Helsinki declaration and its later amendments or comparable ethical standards.

### Microblot array

The collected donors’ sera were analyzed for the presence of multiple antigen-specific antibodies using Microblot-Array COVID-19 IgG, IgA, or IgM kits (TestLine Clinical Diagnostics, Brno, Czech Republic). The analyses were performed according to the manufacturer’s instructions. The levels of the specific antibodies were evaluated according to the manufacturer’s instructions in (U/ml). The samples with (U/ml) values < 185 were negative, between 185–210 borderline, and >210 positive. The IgG, IgA, and IgM antibodies against the following antigens were determined: SARS-CoV-2 spike glycoprotein receptor-binding domain (RBD) and S2 domain (S2), SARS-CoV-2 nucleocapsid protein (NCP), E protein (EP), and papain-like protease (PLP), Middle East respiratory syndrome-related coronavirus (MERS-CoV) spike glycoprotein S1 subunit (S1), SARS-CoV NCP, human coronavirus 229E (HuCoV 229E) NCP, human angiotensin-converting enzyme (ACE-2).

### Peptide-mediated enrichment of the antigen-reactive T Cells

The cryopreserved PBMCs were reconstituted at the concentration 2 × 10^6^ cells/ml in a human plasma serum-containing medium [RPMI 1640 medium, 5% human plasma serum (One Lambda, Canoga Park, CA), 100 U/ml penicillin-streptomycin, 2 mM GlutaMax, 1 mM sodium pyruvate and nonessential amino acid mix (Thermo Scientific)] supplemented with 10 IU/ml of IL-2 (PeproTech, Rocky Hill, NJ, USA). The reconstituted PBMCs were stimulated with 0.5 μg/ml concentration of pooled overlapping peptides spanning the whole molecule of SARS-CoV-2 spike glycoprotein [28] [PepMix™ SARS-CoV-2 Spike Glycoprotein, cat.# PM-WCPV-S-1, JPT Peptide Technologies, Berlin, Germany]. The cells were then cultured for 12 days supplementing the cell cultures with fresh media and IL-2 every 2–4 days. The 12-day cell cultures with the peptide-reactive cells were processed immediately or cryopreserved.

### Cell stimulation and intracellular cytokine staining

The cultured cells or isolated PBMCs were stimulated with 0.5 μg/ml concentration of the peptide pool. After 1 h of culture (37 °C, 5% CO_2_), the cells were supplemented with brefeldin A solution (BioLegend, San Diego, CA) and cultured for 4 h. The samples stimulated with the peptide solvent alone (20% DMSO in PBS) were used as unstimulated controls. The cells were transferred to a V-bottom 96-well plate (Nalgene) and stained as described [29] with live/dead fixable stain and the following antibodies: CD4-PE-Cy7 and CD8-Alexa Fluor 700 (Exbio, Prague, Czech Republic), CD3-PerCP-Cy5.5, TNFα-APC, and IFNγ-PE (Becton Dickinson, Franklin Lakes, NJ). The cells were analyzed by FACSAria II (Becton Dickinson, Heidelberg, Germany) and the data processed by FlowJo software (Tree Star, Ashland, OR). The frequency of reactive T cells was calculated as the difference between the frequency of the cytokine-producing T cells of the vehicle-stimulated sample and the peptide pool-stimulated sample of the same donor.

### Rapid expansion protocol (REP)

The REP was performed for the large-scale expansion of the peptide-enriched cell cultures [17]. As the feeder cells were used PBMCs isolated from buffycoats as described [30]. The buffy coats were obtained from the Institute of Hematology and Blood Transfusion in Prague and each donor provided signed written informed consent to participate in the study. The isolated PBMCs from 3 donors were pooled and γ-irradiated [64 Gy; Gammacell 3000 ELAN (Best Theratronics, Ottawa, ON, Canada)]. The irradiated PBMCs were combined with the peptide-enriched cell cultures, 100 ng/ml of CD3-specific antibody (OKT-3; Miltenyi Biotec, Gladbach, Germany), and cultured in tissue culture flasks (TPP, Trasadingen, Switzerland) for 11 days in the human plasma serum-containing medium with 3000 IU/ml of IL-2 (PeproTech). The cell cultures were supplemented with fresh media and IL-2 every 2–4 days. The expanded cultures were processed immediately or cryopreserved.

### Statistical Analysis

The means of values±SEM were calculated from the indicated sample size (*n*) using GraphPad Prism 6 (GraphPad Software, La Jolla, CA) and the statistical significance (\**p*<0.05, \*\**p*<0.01, \*\*\**p*<0.001, \*\*\*\**p*<0.0001) between two groups of samples determined by Wilcoxon matched-pair signed-rank tests and between three or more groups the statistical significance was determined by matched-pair 1-way ANOVA with Dunn’s post test. The associations between two variables were assessed by the Spearman’s rank-order correlation coefficient (*r*) and the statistical significance of the correlation (*p*) determined. Graphical images were created with Biorender.com (accessed in March and April, 2021).

## Abbreviations

SARS-CoV-2: severe respiratory coronavirus 2
COVID-19: coronavirus disease 2019
RBD: receptor-binding domain
PBMC: peripheral blood mononuclear cell
REP: rapid expansion protocol

## Ethical statement

All experimental protocols were approved by the ethical standards of the institutional and/or national research committee – the Ethics Committee of the University Hospital Motol in Prague, and performed in accordance with the 1964 Helsinki declaration and its later amendments or comparable ethical standards. All patients provided signed informed consent for the use of their blood-derived products for future research.

## Author contributions

P.T., J.L., and Da.S. conducted the experiments and/or analyzed the data; P.T. and Da.S. designed the experiments; Dm.S., Z.S. and J.B. supervised the sample collection and clinical aspects of the study; Da.S. wrote the manuscript; P.T., J.L. Dm.S., Z.S., and J.B. contributed to the writing of the manuscript; Da.S. supervised the research.

## Acknowledgments

We thank the clinical and laboratory research staff for their assistance, Michal Rataj for assistance with the flow cytometry, and John Wilson for his review of the manuscript.

